# Ergonomic risk factors and Work-Related Musculoskeletal Disorders among Fireworks workers in West Bengal, India: A cross sectional study

**DOI:** 10.1101/2021.06.07.447237

**Authors:** Payel Laskar, Subhadeep Ganguly, Zakir Md Hossain

**Author notes:** Address for correspondence /.

## Abstract

**Objective:** Fireworks industries are very old, unorganized cottage industries in West Bengal mainly confined in South 24 Parganas. The present investigation was intended to investigate the prevalence of work-related musculoskeletal disorders among the workers and to identify the causative factors behind it.

**Method:** In this present study, 152 male fireworks workers from different age groups and 100 control subjects were investigated. Modified Nordic questionnaire were used to identify the region wise disorders. Hand Grip strength of both groups were also estimated.

**Result:** Among the fireworks workers posture related musculoskeletal disorders were severely observed in the lower back which was aggravated with the advancement of age and working experience. Pain and stiffness were also reported in neck, upper back, wrist, elbow, knee and ankle. Lower back rest with support at lumber region was strongly recommended.

**Conclusion:** After analysis of ergonomic factors and results, it can be concluded that the fire workers who are working with awkward postures have high risk of developing WMSDs specially affecting the upper limbs and both upper and lower back.

## 1. INTRODUCTION

Fireworks industries are well known ancient but hazardous cottage industries in India. Fireworks mainly emit light, sound, gas and heat on ignition of pyrotechnic chemicals (Cao X et al.,2017). It creates extensive environmental pollution within a short time, deposit metal dust, toxins and other harmful chemicals. Workers involved in the manufacturing process get continuously exposed to it and thus are affected adversely. In West Bengal, the largest fireworks hub is situated in Champahati, South 24 Parganas which includes around 19 villages spread across four gram-panchayats. Approximately 5000 workers are involved in this profession which involves 800 trades directly with an estimated revenue collection of around 25 crores per annum (a report published on The Times of India, October’26, 2019).

Disabilities associated with Work-related Musculoskeletal Disorders (WMSDs) are very common with rapidly increasing socio-economic problem in industrial sectors (Turner JA et al., 2004; Hartvigsen J et al., 2013; Freire ACGF et al., 2017, Hossain MD et al.,2018). Risk factors that aggravate the situation include posture, material handling, mechanical compression, force, temperature, extremities, vibration, glare, duration of exposure, inadequate lighting, etc (Vandergift JL et al., 2010; Comper MLC et al., 2013; Lucas RAI et al., 2014; Golchha V et al., 2014; Charles LE et al., 2018; Noor U et al., 2017; Saedpanah K et al, 2017; Kim JY et al., 2018; Mohan SB 2018;). Many scientists reported that physical burdens and psychological job demand or control also have an impact on musculoskeletal pain.

But very few reports are available on the health hazards associated with the fireworks workers. Like other industrial workers, WMSDs are very common among the fireworks workers. Musculoskeletal disorders are commonly referred to as injuries and disorders to the muscles, tendons, joints, ligaments, cartilage, nerves or spinal disc (Buckle P, 2005). Postural discomforts are associated with works related musculoskeletal disorders mainly in lower back, shoulder, upper extremities among the workers. So far as information available, probably it is the first report of works related musculoskeletal disorders among the fireworks workers in the state of West Bengal. The present work was aimed to assess the ergonomic risk factors associated with the development of WMSDs among male fireworks workers in West Bengal, India.

## 2. METHODS

### 2.1. Study Period & Working Area

The study was conducted from April 2018 to March 2020 involving the fireworks workers of Champahati region of South 24 Parganas district of West Bengal, in India.

### 2.2. Selection of Subjects

The fireworks industry is dominated by male sex workers. Thus, for this study, 152 male fireworks workers were selected for experimental group and 100 workers who are not attached with the field of fireworks and have no involvement in the awkward postures like fireworks workers were selected for control group. The experimental group were divided into four subgroups according their age- A: 18-29years, B: 30-39years, C: 40-49 years, and D: above 50 years. Both groups were physically and mentally healthy. The workers of the control group belonged to different fields (police, shopkeepers, teacher, etc who work at different office and place) other than fireworks who live in that area.

### 2.3. Inclusion and exclusion criteria

Inclusion criteria included age of the workers greater than 18 years and who were involved in the manufacturing process of fireworks for minimum of last 2 years with an average activity of 8 hours on an average per day. All selected workers answered the Nordic questionnaires and participated in the postural analysis. Female workers and physically disabled male workers were excluded from the present study; as female workers in this industry are mostly associated with the packaging department, not affiliated with the manufacturing department.

### 2.4. Physical Parameters

The height and weight of the fireworks workers and the control group were measured with a Martin anthropometer (Takei, Japan) and Crown weighing machine (Raymon Surgical, India) respectively. The body surface area (BSA) was calculated based on Kuorinka I et al.,1987. The BMI (body mass index) of all the subjects were also computed as per Cole TJ et al.,2000.

### 2.5. Questionnaire

A modified Nordic questionnaire was used for this study (Dickinson CE et al.,1992). The questionnaire was represented in the form of multiple choices. The subjects were informed about the objective of the study and all had their consent. The questions were divided into two groups on the base of general information of workers and pain of discomfort body parts.

### 2.6. Statistical Analysis

The mean, standard deviation, median, absolute frequencies and percentages were used to define sociodemographic and other different variables. Student’s t test was used between the two groups of workers to find out whether there was any significant difference between the demographic variable of the experimental and the control groups. Linear correlation and regressions were done to detect the magnitude and direction of association between two variables. P < 0.05 was considered as statistically significant. Most of the statistical analysis were carried out using Microsoft Excel.

### 2.7. Analysis of Working Posture

To analyze the working posture, REBA (rapid entire body assessment) which was proposed by Hignett et al, 2000 to assess posture for risk of work-related musculoskeletal disorders (WRMSDs) was used to analyze the different working posture along with RULA (rapid upper limb assessment) (M Yusuf et al.,2016).

### 2.8. Discomfort Level Scale

To identify the discomfort level, score a ten-point scale was used (S Gangopadhyay et al.,2010). Zero (0) represented ‘no discomfort’ and ten (10) represented ‘worst discomfort (extremely uncomfortable)’. This scale was used to identify the discomfort level of the fireworks workers in their different postures. The intensity of pain or discomfort was measured by utilizing the body part discomfort (BPD) scale.

### 2.9. Risk level scale

After identifying by REBA and RULA method of different postures of fireworks workers, the risk level of the different postures was calculated using a Risk level scale. The Risk Level Scale is also a ten (10) point scale for assessment the level of risk in working posture, where 1 represented ‘no risk or negligible’ and 10 represented ‘very high risk’.

### 2.10. Ethical considerations

All procedures performed in studies involving human participants were in accordance with the ethical standards of 1964 Helsinki declaration and its later amendments. All participants under study signed a written informed consent form. The project was approved by the Institutional Human Ethics Committee of the Aliah University (AU/2022/MZH/IEC003)

## 3. RESULTS

Taken together 252 participants completed the Nordic questionnaire. The overall demographic data of Fireworks workers and control group is presented in Table - 1. There was no significant difference in mean age and BMI between the two groups. However, few differences were observed in the height, weight, BSA and year of experience. Duration of work per day, duration of rest in between work per day and number of working days per week are same for the two groups.

**Table 1:** Sociodemographic data of the fireworks workers and control groups.

|  |  |  |  |
| --- | --- | --- | --- |
| Variable | Fireworks workers (Mean±SD) | Control group (Mean±SD) | p value |
| Age (year) | 37.43 ± 11.81 | 36.10±11.00 | 0.159 |
| Height (cm) | 165.55 ± 8.44 | 168.58±6.68 | 0.008 |
| Weight (Kg) | 59.74 ± 11.45 | 62.32±10.26 | 0.032 |
| BSA (m2) | 1.65 ± 0.17 | 1.71±0.15 | 0.003 |
| BMI (Kg/m2) | 21.80 ± 3.93 | 21.86±2.86 | 0.446 |
| Year of experience | 16.70 ± 10.91 | 11.41±10.58 | 0.0001 |
| Duration of work per day (Hours) | 8 | 8 |  |
| Duration of rest in between work per day (Hour) | 1 | 1 |  |
| Number of working days per week | 6 | 6 |  |

The firework workers are ill-trained and work in a harsh environment without much protective gears and comfort. The manufacturing set up is also improper with too large quantities of explosives and too little distance. Some of their working activities are demonstrated below-

The process of manufacturing of fireworks is a complex process involving multiple process, specialized skills and supervision. It is ready for sale after going through the following six procedures that are depicted below-

All the participant workers’ musculoskeletal problems were evaluated using the Nordic Questionnaire. Everyone who was asked to fill out the survey did so, making up a 100% participation rate. The modified Nordic musculoskeletal questionnaire analysis revealed that discomfort in different body parts among fireworks workers are predominantly reported in knee, lower back, and upper back regions. Discomforts in neck, shoulder, elbow and wrist are also found (Table 2).

**Table 2:** Comparison of discomfort in different body parts between Fireworks workers and control group.

| Age groups | Body parts involved | Fireworks workers | Control group |
| --- | --- | --- | --- |
| Group A |  | N = 44 | N = 28 |
| 20-29 years | Neck | 8 (18.18%) | 5 (17.85%) |
|  | Shoulder | 4 (9.09%) | - |
|  | Elbow | 4 (9.09%) | - |
|  | Wrist | 4 (9.09%) | - |
|  | Upper back | 3 (6.82%) | - |
|  | Lower back | 4 (9.09%) | - |
|  | Hip | - | - |
|  | Knee | - | - |
|  | Ankle | - | - |
| Group B |  | N = 41 | N = 49 |
| 30-39 years | Neck | 11 (26.83%) | - |
|  | Shoulder | 10 (24.39%) | - |
|  | Elbow | 12 (29.27%) | 7 (14.28%) |
|  | Wrist | 12 (29.27%) | - |
|  | Upper back | 21 (51.22%) | 17 (34.69%) |
|  | Lower back | 22 (53.66%) | 19 (38.77%) |
|  | Hip | - | - |
|  | Knee | 8 (19.51%) | - |
|  | Ankle | 1 (2.44%) | - |
| Group C |  | N = 39 | N = 15 |
| 40-49 years | Neck | 11(28.20%) | 2 (13.33%) |
|  | Shoulder | 10(25.64%) | 8 (16.32%) |
|  | Elbow | 14(35.89%) | - |
|  | Wrist | 15(38.46%) | - |
|  | Upper back | 22(56.41%) | 4 (26.66%) |
|  | Lower back | 26(66.67%) | 6 (40.00%) |
|  | Hip | - | - |
|  | Knee | 13(33.33%) | 3 (20.00%) |
|  | Ankle | 6(15.38%) | - |
| Group D |  | N = 28 | N = 8 |
| 50 years and above | Neck | 9(32.14%) | - |
|  | Shoulder | 9(32.14%) | 1 (6.66%) |
|  | Elbow | 10(35.71%) | - |
|  | Wrist | 11(39.29%) | - |
|  | Upper back | 18(64.29%) | 6 (75.00%) |
|  | Lower back | 22(78.57%) | 7 (87.50%) |
|  | Hip | - | - |
|  | Knee | 14(50%) | 7 (87.50%) |
|  | Ankle | 4(14.29%) | - |

The modified Nordic questionnaire also revealed the different injuries associated with the workers. The types of injury in different body parts associated with fireworks are presented in Table-3.

**Table 3.** Types of injury between Fireworks workers and control group.

| **Table 3. Types of injury between Fireworks workers and control group** |  |  |  |  |
| --- | --- | --- | --- | --- |
| Type of injury | Body parts |  | Fireworks workers (%) | Control group (%) |
| Pain | Hand | Elbow | 17.76 | 7 |
|  |  | Wrist | 19.08 | 0 |
|  |  | Finger | 19.73 | 9 |
|  | Leg | Knee | 19.79 | 10 |
|  |  | Ankle | 7.24 | 0 |
|  |  | Feet | 11.18 | 5 |
| Tingling | Hand | Elbow | 1.32 | 0 |
|  |  | Wrist | 3.04 | 0 |
|  |  | Finger | 3.04 | 0 |
|  | Leg | Knee | 1.32 | 0 |
|  |  | Ankle | 0.66 | 0 |
|  |  | Feet | 0.66 | 0 |
| Sprain | Hand | Elbow | 5.26 | 0 |
|  |  | Wrist | 11.84 | 0 |
|  |  | Finger | 16.44 | 0 |
|  | Leg | Knee | 0.66 | 0 |
|  |  | Ankle | 14.47 | 3 |
|  |  | Feet | 13.16 | 0 |
| Abrasion | Hand | Elbow | 3.29 | 0 |
|  |  | Wrist | 16.45 | 1 |
|  |  | Finger | 16.45 | 1 |
|  | Leg | Knee | 1.32 | 0 |
|  |  | Ankle | 4.6 | 0 |
|  |  | Feet | 5.26 | 0 |
| Numbness | Hand | Elbow | 4.6 | 0 |
|  |  | Wrist | 5.92 | 0 |
|  |  | Finger | 6.58 | 0 |
|  | Leg | Knee | 4.6 | 0 |
|  |  | Ankle | 3.29 | 0 |
|  |  | Feet | 2.63 | 0 |
| Swelling | Hand | Elbow | 1.32 | 0 |
|  |  | Wrist | 7.9 | 2 |
|  |  | Finger | 8.6 | 3 |
|  | Leg | Knee | 1.97 | 0 |
|  |  | Ankle | 1.32 | 0 |
|  |  | Feet | 6.6 | 4 |
| Laceration | Hand | Elbow | 3.29 | 0 |
|  |  | Wrist | 7.89 | 0 |
|  |  | Finger | 10.53 | 2 |
|  | Leg | Knee | 1.97 | 0 |
|  |  | Ankle | 2.63 | 0 |
|  |  | Feet | 11.84 | 1 |

REBA offers a quick and simple way to evaluate the risk of WMSDs associated with various working postures. It provides a scoring system for muscle activity over the complete body and separates the body into portions to be independently coded in accordance with movement planes. RULA assesses the strength and activity of the muscles that lead to repetitive strain injuries (bad posture or attitude). The risk level of the different postures using REBA and RULA was analyzed as a ten (10) point scale which are described in Table 4.

**Table 4.**
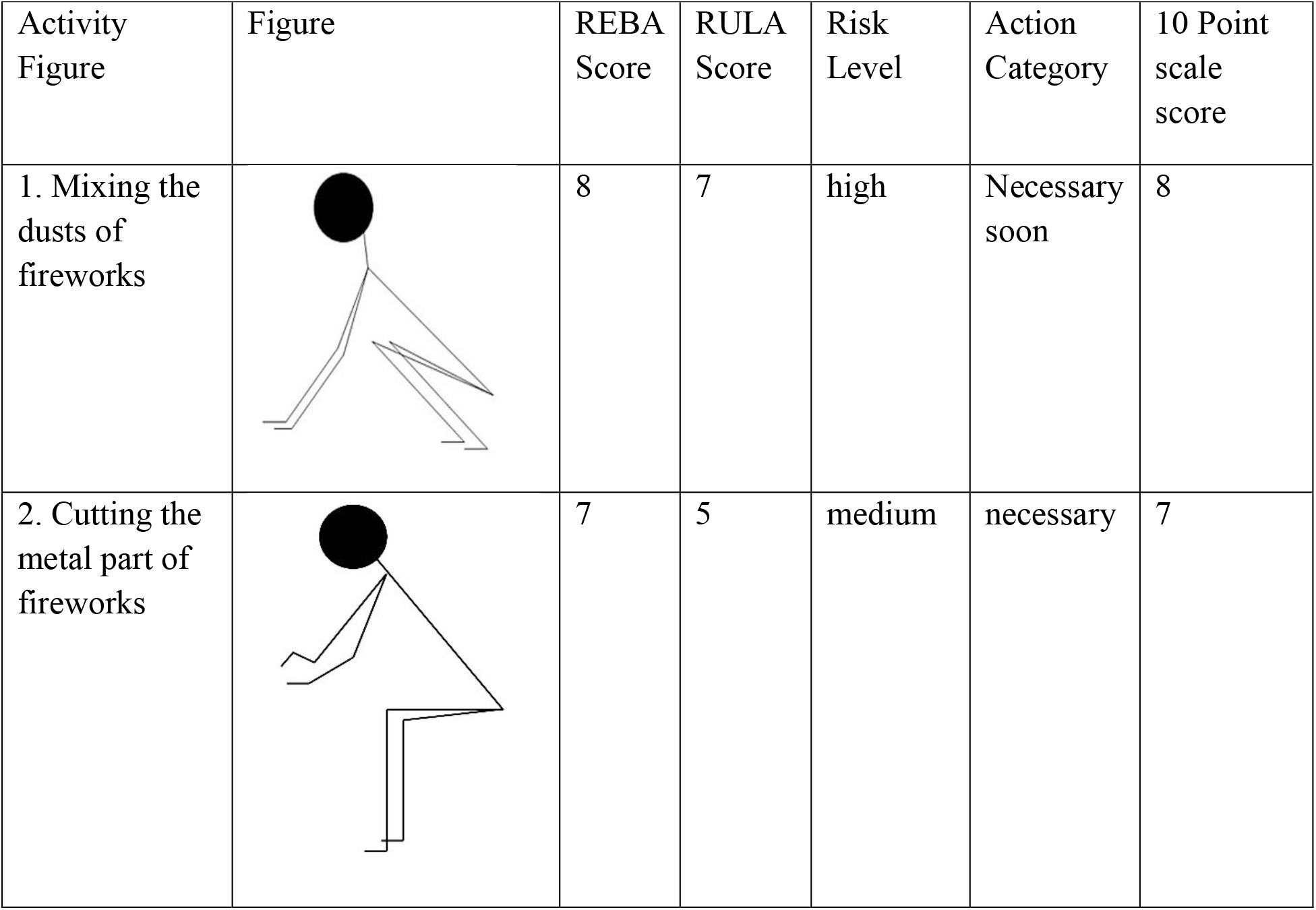
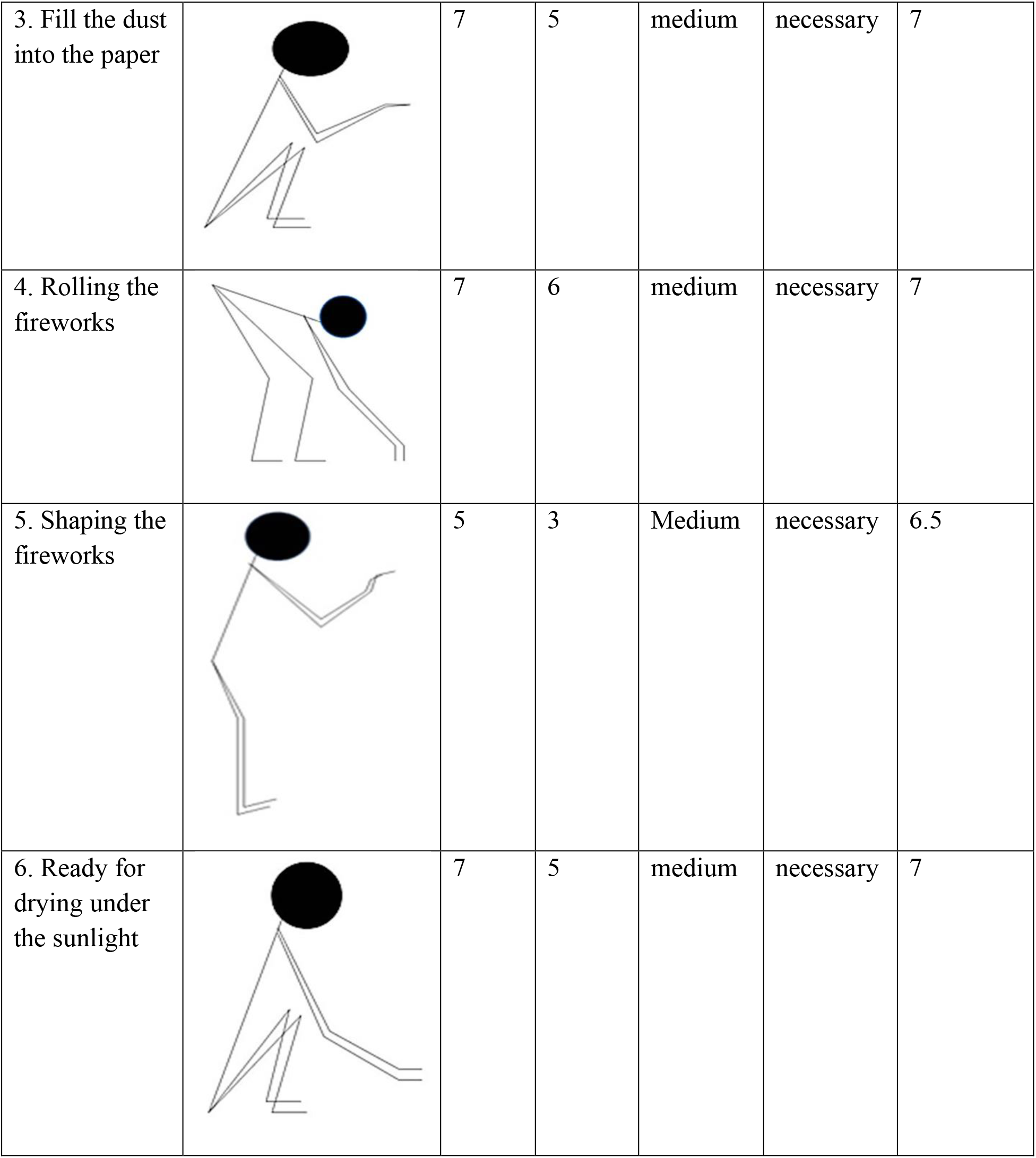
Analysis of working posture of Fireworks workers by REBA and RULA.

| Activity Figure | Figure | REBA Score | RULA Score | Risk Level | Action Category | 10 Point scale score |
| --- | --- | --- | --- | --- | --- | --- |
| 1. Mixing the dusts of fireworks |  | 8 | 7 | high | Necessary soon | 8 |
| 2. Cutting the metal part of fireworks |  | 7 | 5 | medium | necessary | 7 |
| 3. Fill the dust into the paper |  | 7 | 5 | medium | necessary | 7 |
| 4. Rolling the fireworks |  | 7 | 6 | medium | necessary | 7 |
| 5. Shaping the fireworks |  | 5 | 3 | Medium | necessary | 6.5 |
| 6. Ready for drying under the sunlight |  | 7 | 5 | medium | necessary | 7 |

Linear regression between discomfort levels and risk levels at different working posture of the entire body parts were calculated which is depicted in Figure 3a, whereas Figure 3b shows the discomfort found at different times among fireworks workers (%) and control group (%). The correlation between the entire body parts and discomfort level is shown in Table 5. The lower body parts show significant (p<0.05) positive correlation (r=0.862) with discomfort level.

**Fig.1-.**
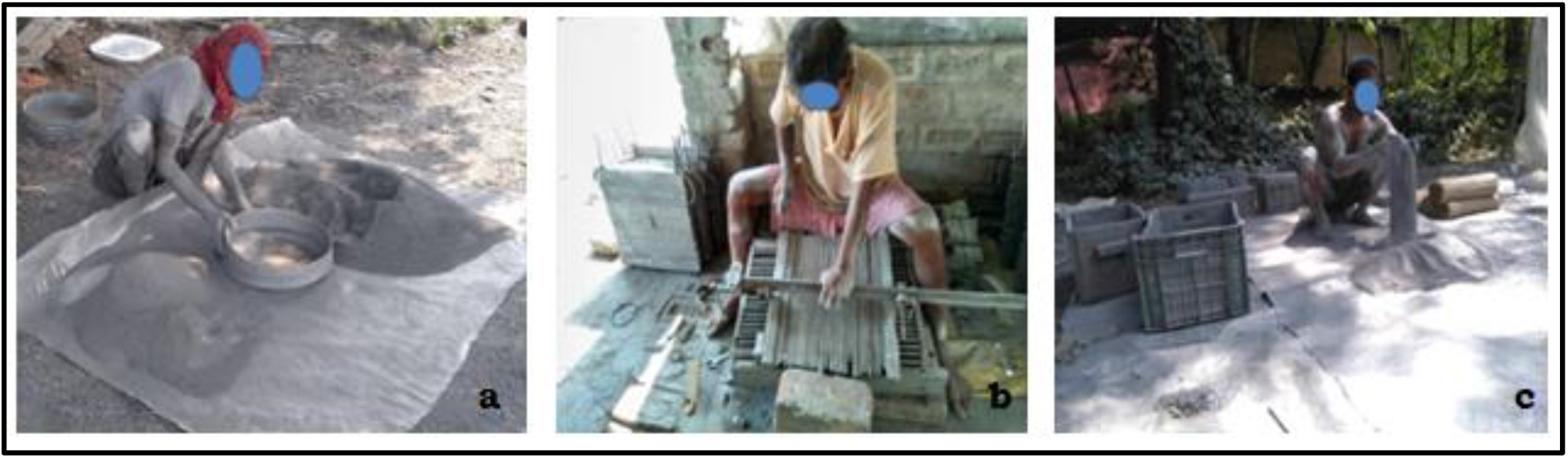
(a) Mixing the dusts of fireworks (b) Cutting the metal part (c)Fill the dust into the paper

**Fig.2-.**
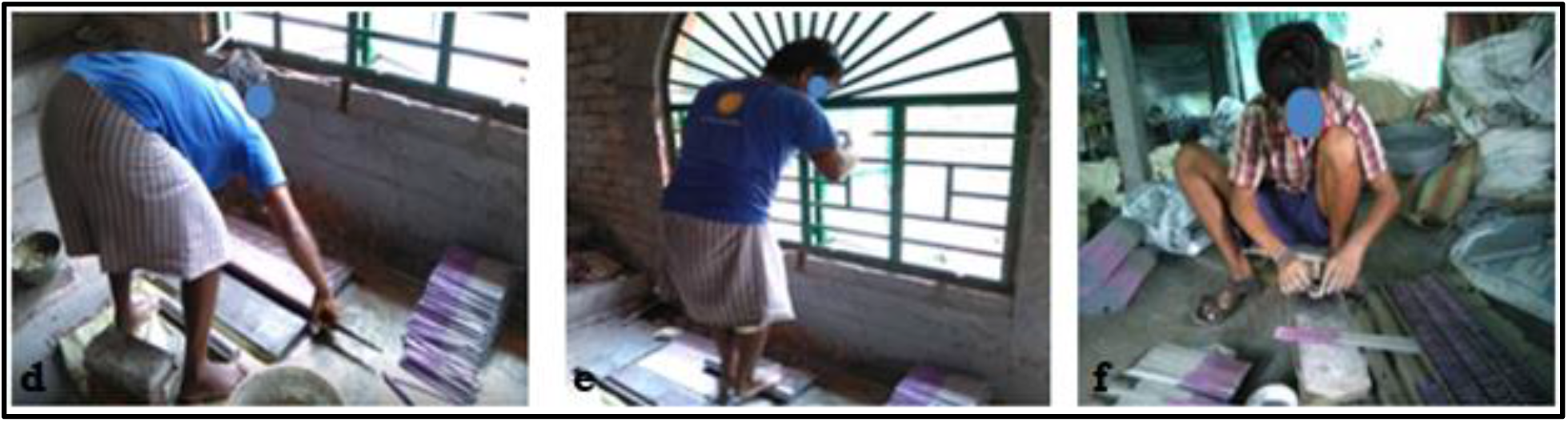
(d) Rolling the fireworks (e) Shaping the fireworks (f) Ready for drying in the sunlight

**Fig.3:**
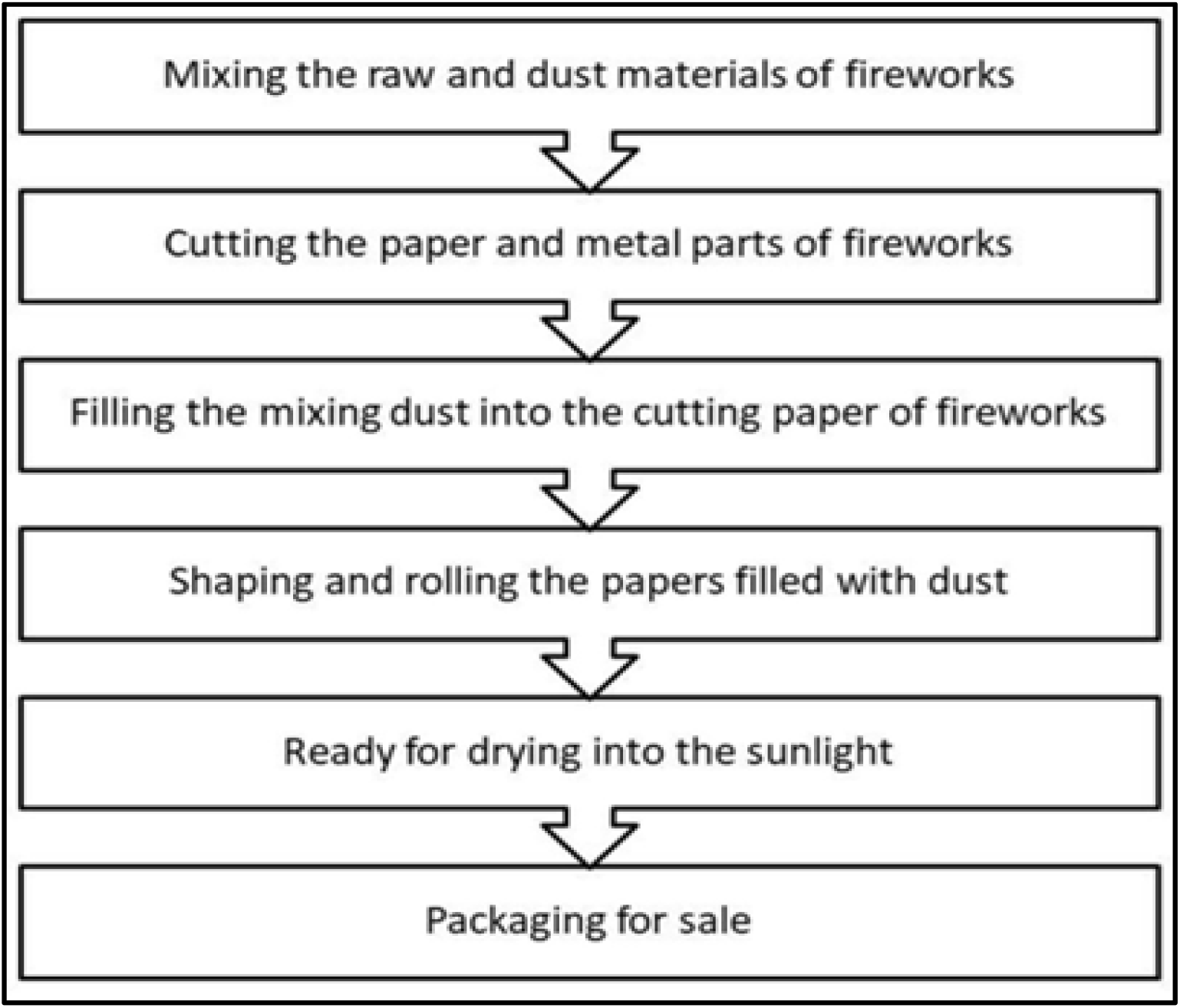
Flow chart of fireworks making process

**Fig.3a (left):**
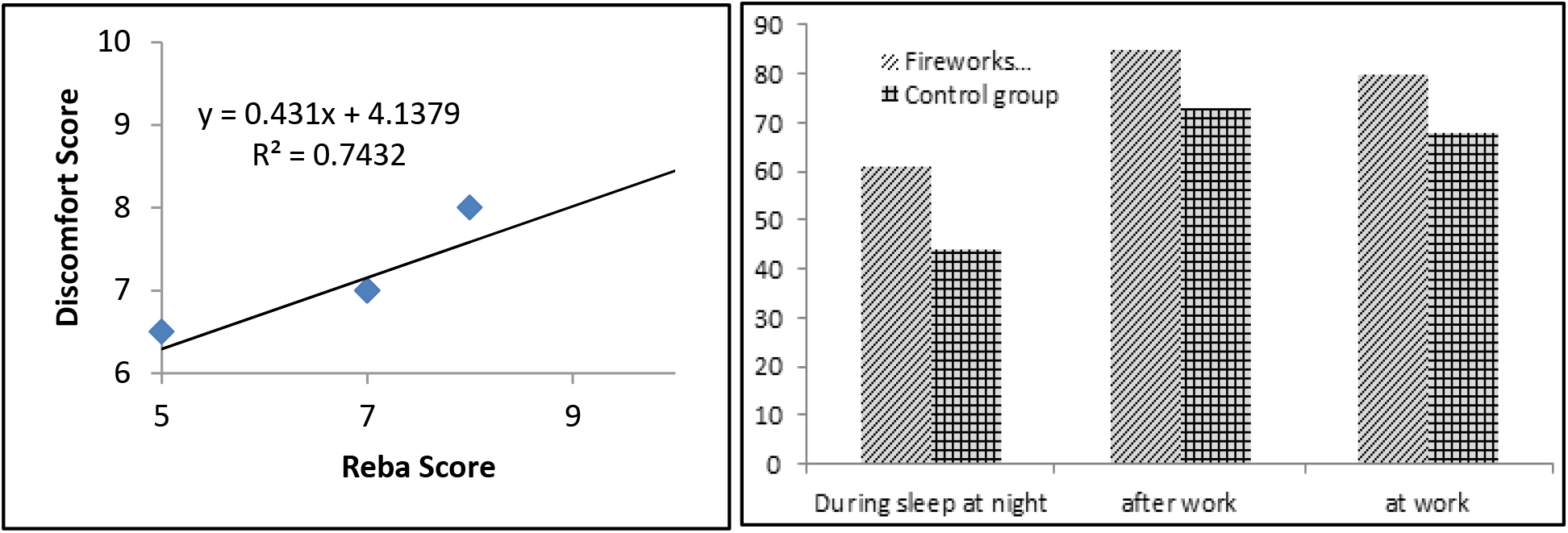
Linear regression between discomfort level and risk level at entire body parts in different working posture; Fig.3b (right): Discomfort at different times among fireworks workers (%) and control group (%)

**Table 5.**
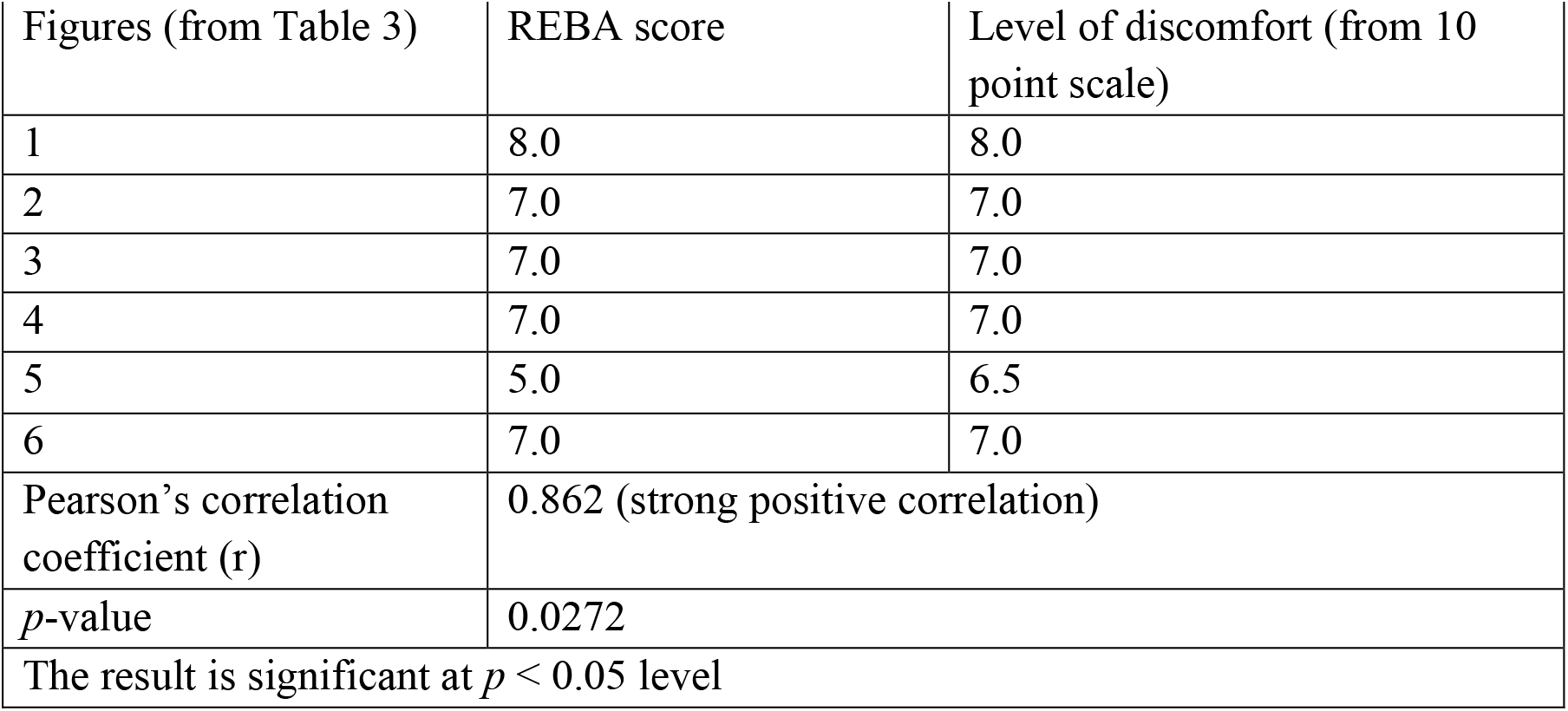
Correlation between entire body parts and discomfort levels.

| Table 5. Correlation between entire body parts and discomfort levels |  |  |
| --- | --- | --- |
| Figures (from Table 3) | REBA score | Level of discomfort (from 10 point scale) |
| 1 | 8.0 | 8.0 |
| 2 | 7.0 | 7.0 |
| 3 | 7.0 | 7.0 |
| 4 | 7.0 | 7.0 |
| 5 | 5.0 | 6.5 |
| 6 | 7.0 | 7.0 |
| Pearson's correlation coefficient (r) | 0.862 (strong positive correlation) |  |
| p-value | 0.0272 |  |
| The result is significant at $p < 0.05$ level | | |

## 4. DISCUSSION

Work related musculoskeletal disorders are rapidly increasing among workers working in both large-scale industries as well as in small scale cottage industries in India. Various socio-economic conditions like poverty associated with malnutrition, unscientific working posture, adverse working conditions, excessive working hours, etc aggravate the situation. It is evident that fireworks workers working in unfavorable conditions might have developed some work-related musculoskeletal disorders. From the observation of working condition of fireworks workers, it can be said that they spent their maximum times of day with toxic chemicals and dusts of chemicals. They dry the fireworks under the sunlight, doing so they face very hot temperature of the sun.

They complete their work by sitting with awkward posture which might be the reason that they have maximum discomfort in lower back (48.7%). The details are shown in Table-2. Pain of Upper back (42.1), Knee (23%) also are seen which may be for the awkward sitting procedure. They are involved in working with fine skilled activity like packaging fireworks that requires them to bend their head and neck. Maybe that’s the reason it causes discomfort of their neck (25.75), shoulder (21.7%), elbow (26.3%), wrist (27.6%). The intensity and frequency of pain and different type of discomfort increase with time.

Shyam and Dutt (2017) reported significant relation between age, years and working hours as well as marital status with the development of work-related musculoskeletal disorders which affect almost 70% of the working group involved in textile industries of Meerat. Prevalence of work-related musculoskeletal disorders with 58.8% of coir industry workers were also reported in Kerala. The same thing is seen in our present study on fireworks workers. Increased prevalence of work-related musculoskeletal disorder is seen with advancement of age. In Table-2, it is clearly seen that maximum workers affected with MSD are in the age group of D (50 and above) compared to other or younger age groups. Maximum work-related musculoskeletal disorders involving the lower back were found maximally up to 71.73% fireworks workers in Group C (age group 40-49 years). However, pain and stiffness were also reported in upper back, neck, wrist, elbow, knee and ankle.

Studies showed that WRMSD is caused by awkward working posture, repetitive, forceful activities and carrying overload. Analysis of the working posture of fireworks workers which was done by the REBA and RULA method is described in Table 4. REBA and RULA scores reveal that all postures needed to be rectified ergonomically. The posture of mixing the dust is the most harmful among the all postures. They work repetitively with this working posture throughout the day. Maybe these are the reasons for the development of MSD among fireworks workers (Marta et al.,2021, Lincoln et al.,2022). The postures which are performed more than 4 times in one minute increase the score of RULA (INERMAP, 2011; McAtamney and Corlett, 1993). Cutting the metal part of fireworks, filling the dust into the paper, rolling the fireworks, and shaping the fireworks are those postures that are performed more than 4 times in one minute. Figure 3a shows the linear regression between discomfort level and risk level at entire body parts in the different working postures with the value of R2 = 0.7432. The correlation between the entire body parts and discomfort level is shown in Table 5. The lower body parts show a significant (p<0.05) positive correlation (r=0.862) with discomfort level.

Chaffin and Anderson and Leskinen said that the number of forward bend involved in working posture influence the compressive strength of erector spine muscle and vertebral disc. The present study has shown that fireworks workers sit with a forward bend posture in fireworks making and packaging for a prolonged time. Maybe this is another reason for developing lower back pain among the experimental group as shown in Table 3. This investigation also has shown that fireworks workers said about their discomfort after work. During work they also feel discomfort. But they are not too much conscious about their body parts discomfort. Figure 4 has shown the consciousness of fireworks workers about their discomfort body parts. Very few workers said about their discomfort during sleep. Some workers complain about sleep disturbance due to discomfort of body parts. The intensity of discomforts of body parts among fireworks workers is higher than control group as depicted in Fig. 3b.

**Fig. – 4.**
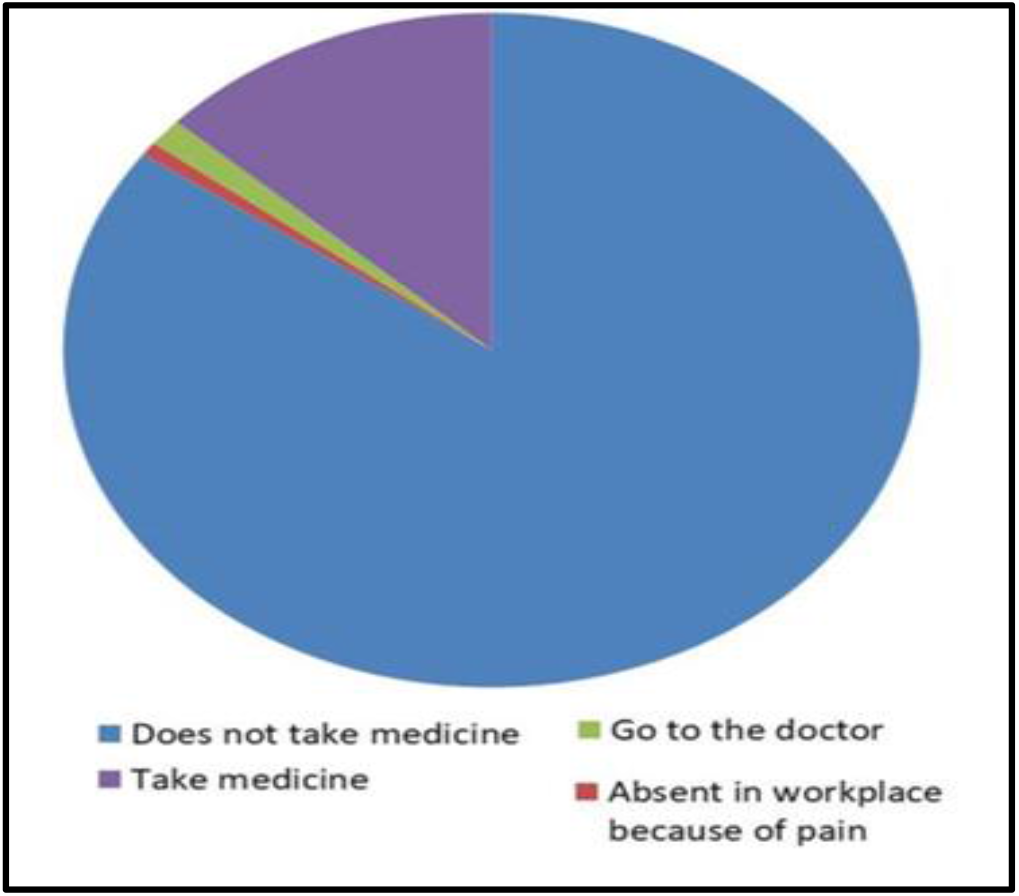
Consciousness of fireworks workers about their discomfort body parts

## 5. CONCLUSION

After analysis of ergonomic factors and results, it can be concluded that the fire workers who are working with awkward postures have high risk of developing WMSDs specially affects the upper limbs and both upper and lower back. Unfavorable working environment, unscientific working postures with psychosocial factors and age significantly caused the development of local and multisite symptoms. Some ergonomics equipment’s are recommended to reduce musculoskeletal discomfort in the lower back, upper back, neck, shoulder and knee of fireworks workers. Different types of exercises, such as stretching etc., are also advised to be practiced for fireworks workers to ease the discomfort.

## 6. FUTURE RESEARCH

The present study had some limitations. To identify long term biochemical and physiological stress among fireworks workers have to done a retrospective study. An electromyographic study and x-ray of discomfort body parts have to be performed. Also need to study of known comorbidities, such as thyroid, hypertension, diabetes etc, that promote to musculoskeletal disorders.

## 7. ACKNOWLEDGEMENT

The authors would like to acknowledge and thank the fire workers of Champahati who kindly participated in this research along with all others who provided their help in any form. The authors also acknowledge Govt of West Bengal for the fellowship provided to Payel Laskar.

